# Epigenetic age-predictions in mice using pyrosequencing, droplet digital PCR or barcoded bisulfite amplicon sequencing

**DOI:** 10.1101/2020.07.30.228122

**Authors:** Yang Han, Miloš Nikolić, Michael Gobs, Julia Franzen, Gerald de Haan, Hartmut Geiger, Wolfgang Wagner

## Abstract

Age-associated DNA methylation reflects aspects of biological aging - therefore epigenetic clocks for mice can help to elucidate the impact of treatments or genetic background on the aging process in this model organism. Initially, age-predictors for mice were trained on genome-wide DNA methylation profiles, whereas we have recently described a targeted assay based on pyrosequencing of DNA methylation at only three CG dinucleotides (CpGs). Here, we have re-evaluated pyrosequencing approaches in comparison to droplet digital PCR (ddPCR) and barcoded bisulfite amplicon sequencing (BBA-seq). At individual CpGs the correlation of DNA methylation with chronological age was slightly higher for pyrosequencing and ddPCR as compared to BBA-seq. On the other hand, BBA-seq revealed that neighboring CpGs tend to be stochastically modified in murine age-associated regions. Furthermore, the binary sequel of methylated and non-methylated CpGs in individual reads can be used for single-read predictions, which may reflect heterogeneity in epigenetic aging. In comparison to C57BL/6 mice the epigenetic age-predictions using BBA-seq were also accelerated in the shorter-lived DBA/2 mice, and in C57BL/6 mice with a lifespan quantitative trait locus of DBA/2 mice. Taken together, we describe further optimized and alternative targeted methods to determine epigenetic clocks in mice.

## Introduction

Aging evokes dynamic changes in DNA methylation (DNAm) at specific CG dinucleotides (CpG) (Dor & Cedar, 2018). These epigenetic modifications provide a biomarker for the aging process, which is often referred to as ‘epigenetic clock’ (Horvath & Raj, 2018). They were initially described for humans based on data from Illumina BeadChips (Bocklandt et al., 2011; Koch & Wagner, 2011), and in the advent of a fast growing number of such datasets the models were further refined - with signatures of many age-associated CpGs - to provide a very high correlation of predicted and chronological age. Notably, epigenetic clocks for blood seem to reflect aspects of biological age, since the deviation of predicted and chronological age (delta-age) correlates with all-cause mortality (Belsky et al., 2020; Marioni et al., 2015) and it is increased in various diseases, such as obesity (Horvath et al., 2014), Down syndrome (Horvath et al., 2015), Werner Syndrome (Maierhofer et al., 2017), and HIV infection (Gross et al., 2016). Thus, tracking of epigenetic age may also elucidate the impact of drugs or other relevant parameters for the aging process, albeit it is challenging to perform such controlled and long-term aging intervention studies in humans (Fahy et al., 2019).

Mice are one of the most popular mammalian models for aging research. Inbreeding, defined growth conditions, and the shorter life span of about two years facilitate aging interventions studies with mice that cannot be easily performed in humans. Epigenetic clocks for mice were initially based on whole genome bisulfite sequencing (WGBS) or reduced representation bisulfite sequencing (RRBS) (Wagner, 2017). They were trained for liver, whole blood, or even multi-tissue specimens from mice using hundreds of CpG sites, and they clearly demonstrated that epigenetic clocks in mice are affected by genetic, dietary, or pharmacological interventions (Petkovich et al., 2017; Stubbs et al., 2017; Wang et al., 2017). However, WGBS and RRBS are relatively labor and cost-intensive and the methods do not always provide enough coverage for all the relevant CpGs, which hampers application of these age-predictors.

To overcome these problems, alternative methods for site-specific analysis of DNAm at few selected age-associated CpGs may be advantageous (Maegawa et al., 2017; Wagner, 2017). We have recently described an epigenetic clock that is based on pyrosequencing of DNAm at only three age-associated CpGs to facilitate a high accuracy with chronological age in C57BL/6 mice (Han et al., 2018). Notably, epigenetic aging was significantly accelerated in the shorter-lived DBA/2 mice (Han et al., 2018), and in congenic C57BL/6 mice harboring regions of chromosome 11 from DBA/2 mice that is likely linked to the regulation of lifespan (referred to as Line A mice) (Brown et al., 2019). The epigenetic age was also decelerated by systemic administration of a drug that extended murine lifespan (Florian et al., Accepted for publication), implying that the three CpGs might also serve as biomarkers of aging at least on an C57BL/6 background. While the pyrosequencing based epigenetic clock has proven to be robust and reliable, it is well conceivable that precision, accuracy and applicability can be increased by alternative methods.

Droplet Digital PCR (ddPCR) is a relatively novel targeted approach for DNAm measurement that was reported to provide precise results with less PCR bias (Han et al., 2020; Zemmour et al., 2018). Furthermore, barcoded bisulfite amplicon sequencing (BBA-seq), which is based on massive-parallel-sequencing, facilitates DNAm analysis of longer amplicons with more neighboring CpGs and provides insight into the DNAm pattern on individual DNA strands (Franzen et al., 2017). We have recently demonstrated in BBA-seq data of human blood that the correlation of age with DNAm levels at neighboring CpGs follows a bell-shaped curved (Han et al., 2020). Interestingly, the DNAm pattern of neighboring CpGs was not coherently modified on individual strands, as might be anticipated upon binding of an epigenetic writer, but rather seemed to be evoked by stochastic modifications (Han et al., 2020). Based on this, we developed an epigenetic age-predictor for BBA-seq data of human blood, which was based on the binary sequel of methylated and non-methylated sites in individual reads (Han et al., 2020). This approach might reflect heterogeneity of epigenetic aging within a sample. In this study, we now established and compared such targeted epigenetic clocks also for mice, which are based on pyrosequencing, ddPCR, BBA-seq, or single read predictions.

## Results

### Alternative epigenetic clocks based on pyrosequencing

In our previous work, we selected nine age-associated genomic regions, which were initially identified for age-predictors based on genome-wide deep-sequencing of DNAm profiles (Petkovich et al., 2017; Stubbs et al., 2017). Based on this, we established a 3 CpG model for pyrosequencing measurements in the genes proline rich membrane anchor 1 (*Prima1*), heat shock transcription factor 4 (*Hsf4*) and potassium voltage-gated channel modifier subfamily S member 1 (*Kcns1*) (Han et al., 2018). Age-predictions correlated very well with the chronological age of C57BL/6 mice in a training set (n = 24; R^2^ = 0.96; Median error = 3.6 weeks) and in two independent validation sets (n = 21 and 19; R^2^ = 0.95 and 0.91; Median error = 5.0 and 5.9 weeks, respectively). We initially also described a 15 CpG model, which considered two additional amplicons of the pseudogene *Gm9312* and myoblast fusion factor (*Gm7325)* (Han et al., 2018). This 15 CpGs model was identified by machine learning and although it provided higher accuracy in the training set (R^2^ = 0.99; Median error = 2.4 weeks), this model was not further validated as we anticipated that the very good correlation might rather be due to overfitting (Han et al., 2018). In present study we further explored this 15 CpG model by pyrosequencing for the two independent validation sets of C57BL/6 mice (n = 21 and n = 19). In fact, 15 CpG clock gave slightly better correlation with chronological age and lower prediction error (R^2^=0.97 and R^2^=0.95; median error = 4.9 weeks and 5.4 weeks, respectively) than the 3 CpG signature (Figure 1). Thus, the 15 CpG murine epigenetic aging clock seems to be advantageous, while the need of two additional PCR amplicons and pyrosequencing measurements provides a tradeoff between accuracy and costs.

**Figure 1.**
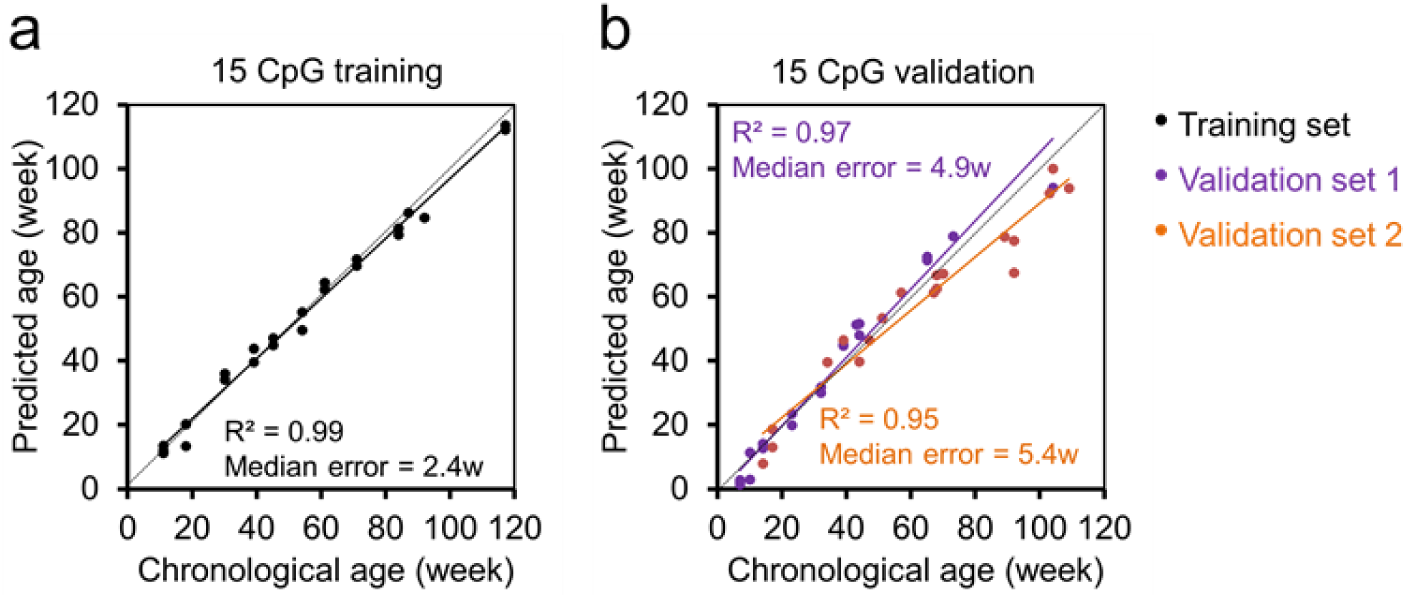
Epigenetic age predictions for pyrosequencing data (15 CpG lasso regression model). (**a**) Multivariable machine learning (Lasso regression) age-predictor based on DNAm levels at 15 CpGs in the genes *Prima1, Hsf4, Kcns1, Gm9312, Gm7325.* Pyrosequencing was performed for 24 C57BL/6 mice (training set) as described before (Han et al., 2018). (**b**) Age predictions with the same model in two independent validation sets: 21 C57BL/6 mice from the University of Ulm and 19 C57BL/6 mice from the University of Groningen (validation sets 1 and 2, respectively). Coefficients of determination (R^2^) of DNAm *versus* chronological age and median errors (weeks) are indicated.

### Age-prediction with droplet digital PCR

Droplet digital PCR (ddPCR) is based on parallel PCR reactions in thousands of micro-droplets and therefore DNAm analysis with this technology may reduce PCR bias for methylated/non-methylated strands that may occur in pyrosequencing (Figure S1a) (Han et al., 2020). Therefore, we have designed ddPCR assay for the same three amplicons for *Prima1, Hsf4*, and *Kcns1*. However, the targeted CpG within the *Hsf4* amplicon was different to the pyrosequencing based 3 CpG predictor, as this was better suitable for the ddPCR probe. DNAm measurements with ddPCR at all three CpGs revealed high correlation with chronological age in 23 C57BL/6 mice of the training set (Figure 2a-c), and correlated with the DNAm measurements by pyrosequencing (Figure S1b). Based on the ddPCR measurements we determined a multivariable linear regression model that provided reliable age-predictions in the validation sets (R^2^ = 0.97 and 0.88; median error 5.1 and 7.1 weeks). These results were slightly less accurate than for the 3 CpG clock by pyrosequencing (Figure 2d), which might be due to lower age-association in the neighbouring CpGs of *Hsf4.* Either way, the results demonstrate that DNAm measurements with ddPCR are also well suited for epigenetic clocks in mice.

**Figure 2.**
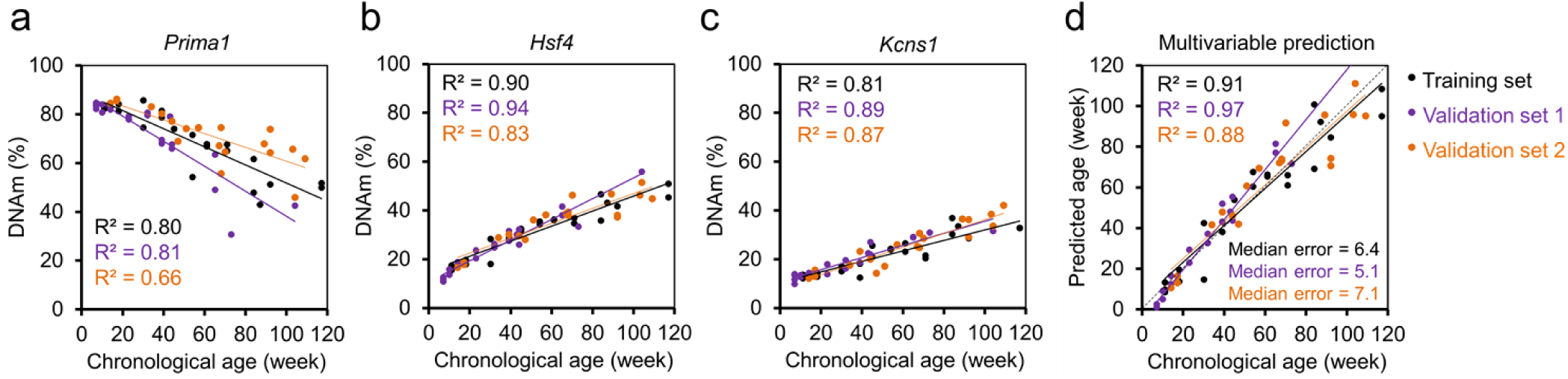
Three CpG epigenetic clock for mice based on droplet digital PCR. Age-associated DNAm was measured with ddPCR at 3 CpGs in the genes *Prima1* (**a**), *Hsf4* (**b**) and *Kcns1* (**c**) in the training set (n = 23) and two independent validation sets (n = 21 and 19) of C57BL/6 mice. (**d**) The measurements of the training set were used for a multivariable model for epigenetic age predictions. Coefficients of determination (R^2^) of DNAm *versus* chronological age and median errors (weeks) are demonstrated.

### Barcoded bisulfite amplicon sequencing of age-associated regions

Subsequently, we used barcoded bisulfite amplicon sequencing (BBA-seq) to investigate age-associated DNAm in amplicons of *Prima1, Hsf4* and *Kcns1*, which covered 4, 12, and 21 neighboring CpGs, respectively. Overall, DNAm measurements correlated in BBA-seq *versus* pyrosequencing (Figure S2), albeit slightly less than ddPCR *versus* pyrosequencing (Figure S1b). Furthermore, the correlation at individual CpGs with chronological age was slightly lower in BBA-seq as compared to pyrosequencing or ddPCR (Table 1). Either way, the three relevant or neighboring CpGs of the pyrosequencing clock also provided a high correlation with chronological age (Figure 3a-c). The BBA-seq measurements of these three CpGs were then used to train a multivariable linear model and the age-predictions correlated well in the validation sets 1 and 2 (n = 21 and 19; R^2^ = 0.95 and 0.91; median error = 6.6 and 10 weeks; Figure 3d). Alternatively, we considered all CpGs of the three amplicons to generate a Lasso regression model with 10-fold cross-validation that considered 7 CpG sites of the three amplicons. The accuracy of age-predictions with this machine learning based model were slightly better for the validation sets (n = 21 and 19; R^2^ = 0.91 and 0.90; median error = 6.1 and 5.9; Figure 3e). Taken together, BBA-seq provided similar accuracy in epigenetic age-predictions as pyrosequencing and ddPCR.

**Table 1.** Correlation of DNAm and chronological age in different targeted approaches.

|  |  | *Prima1* | *Hsf4*<sup>1</sup> | *Kcns1*<sup>1</sup> | Mean R2 |
| --- | --- | --- | --- | --- | --- |
| **Pyrosequencing** | Training ($n = 24$) | 0.71 | 0.96 | 0.84 | 0.84 |
| | Validation1 ($n = 19$) | 0.79 | 0.96 | 0.81 | 0.85 |
| | Validation2 ($n = 21$) | 0.69 | 0.89 | 0.86 | 0.81 |
| **ddPCR** | Training ($n = 23$) | 0.8 | 0.9 | 0.81 | 0.84 |
| | Validation1 ($n = 19$) | 0.81 | 0.94 | 0.89 | 0.88 |
| | Validation2 ($n = 21$) | 0.66 | 0.83 | 0.87 | 0.79 |
| **BBA-seq** | Training ($n = 23$) | 0.78 | 0.87 | 0.85 | 0.83 |
| | Validation1 ($n = 19$) | 0.78 | 0.91 | 0.78 | 0.82 |
| | Validation2 ($n = 21$) | 0.64 | 0.75 | 0.85 | 0.75 |

<sup>1</sup>CpGs from the amplicons were always selected by the highest Pearson correlation with chronological age, therefore they are not identical in the different sequencing approaches.

**Figure 3.**
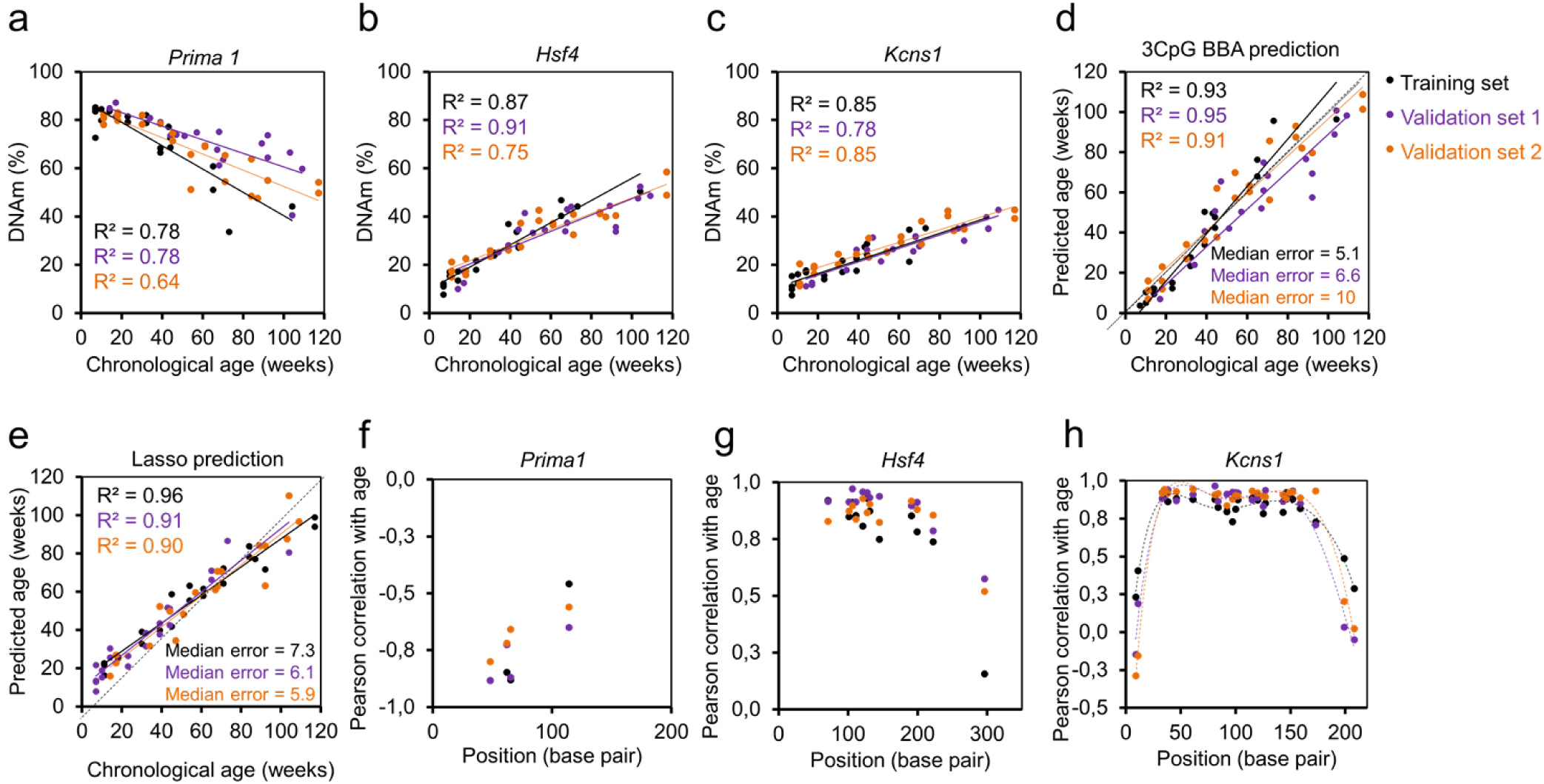
Epigenetic age-prediction by BBA-seq. DNAm levels (%) of three highly age-associated CpGs within three amplicons *Prima1* (**a**), *Hsf4* (**b**) and *Kcns1* (**c**) were determined by barcoded bisulfite amplicon sequencing (BBA-seq). (**d**) Age predictions based on the multivariable linear regression model of three CpGs in the C57BL/6 mice. **(e**) Age predictions with a lasso regression model (7 CpGs in the three age-associated regions), which was trained on the training set of C57BL/6 mice. Coefficients of determination (R^2^) of DNAm *versus* chronological age and median errors (weeks) are indicated. (**f-h**) Pearson’s correlations of age with DNAm levels of CpGs within the amplicons of *Prima1, Hsf4*, and *Kcns1* are plotted for the blood samples of the training set (n = 23) and two independent validation sets (n = 21 and 19). The x-axis represents the position of CpGs within the amplicons.

Subsequently, we analyzed how DNAm at neighboring CpGs correlates with chronological age. For each CpG within the BBA-seq amplicons of *Prima1, Hsf4* and *Kcns1* we determined the correlation with chronological age in the training and validation sets (Figure 3f-h). This analysis revealed that not only the individual CpGs of our age predictor are age-associated, but also the CpGs in the direct vicinity, which is in line with our recent analysis in humans (Han et al., 2020).

### Epigenetic age predictions for mice based on individual BBA sequencing reads

In contrast to pyrosequencing or ddPCR, BBA-seq provides individual reads with a binary sequel of either methylated or non-methylated CpGs. Heatmaps of DNAm within individual reads indicated that the methylation at neighboring CpGs occurs rather independent of each other (Figure 4a and S3a). In fact, Pearson’s correlation of DNAm levels between neighboring CpG sites within the three amplicons revealed only moderate correlation in epigenetic modifications (Figure 4b and S3b), albeit it was slightly higher than previously observed for BBA-seq data in three human age-associated regions (Han et al., 2020).

**Figure 4.**
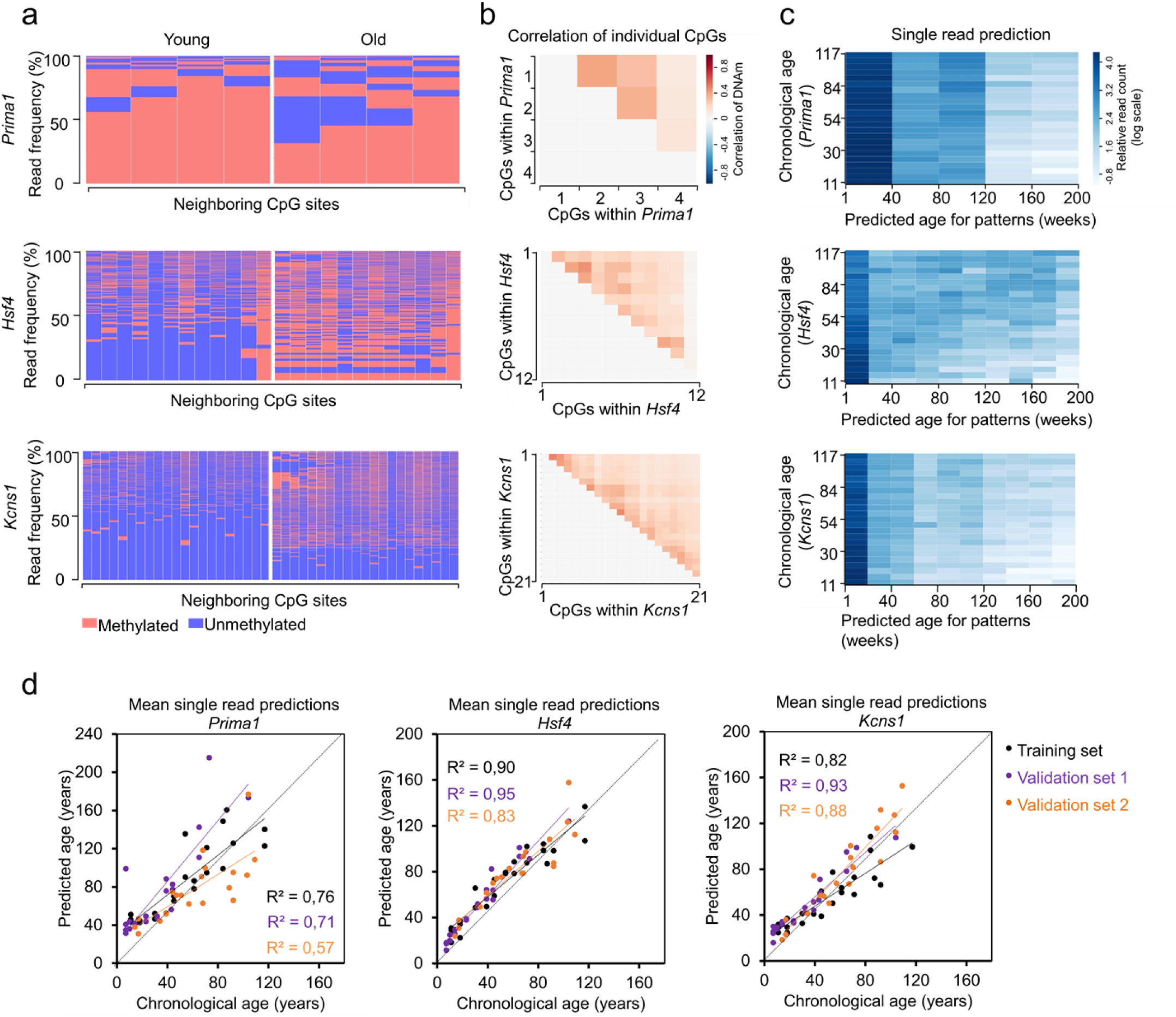
Analysis of age-associated DNAm patterns within individual BBA-seq reads in C57BL/6 mice. (**a**) Frequencies of DNAm patterns in BBA-seq reads (red: methylated; blue: non-methylated) within the amplicons of *Prima1* (4 neighboring CpGs), *Hsf4* (12 neighboring CpGs) and *Kcns1* (21 neighboring CpGs). Samples of one young (11 weeks) and one old C57BL/6 mouse (117 weeks) from the training set are exemplarily depicted. (**b**) Pearson correlation of DNAm among neighboring CpGs within each of the three amplicons in BBA- seq data of the training set. (**c**) Epigenetic ages were estimated for each individual read of the BBA-seq data the training set (n = 23). These single-read predictions were performed for each amplicon based on the binary sequel of methylated and non-methylated CpGs. The heatmaps depict the relative frequency of reads (normalized by the read counts per sample; log scale) that are classified to a specific age category (between 0 and 200 weeks) for each donor in the training set. (**d**) The mean age-predictions based on individual BBA-seq reads of three amplicons were determined for each sample and then plotted against the chronological age of the training (n = 23) and two validation sets (n = 21 and n = 19) of C57BL/6 mice.

For human BBA-seq data we have recently demonstrated that it is possible to estimate the epigenetic age for individual reads, under the assumption that the age-associated modification of DNAm occurs independently at neighboring CpGs. The mean of all individual read-predictions within a sample correlated with the chronological age (Han et al. 2020). Here, we have analyzed if this was also applicable for murine BBA-seq data. For each BBA-seq read of the three amplicons (*Prima1, Hsf4* and *Kcns1*) we estimated the epigenetic age based on the binary sequel of methylated and non-methylated CpGs, using the age-associated correlations at individual CpGs of the training set. Individual reads were predicted between 0 and 200 weeks (Figure 4c and S3c), which might resemble heterogeneity in epigenetic aging within a given sample. Overall the ‘young’ reads were more frequent in young donors, whereas ‘old’ reads were more frequent in old mice. Notably, the mean of single-read predictions within a sample correlated for all three amplicons with the chronological age of the mice (Figure 4d). Particularly for the amplicons of *Hsf4* and *Kcns1*, which harbor more neighboring CpGs, the mean of single read- predictions correlated good or even better than the DNAm levels at the individual age-associated CpGs (Table 1). Thus, it is possible to estimate the epigenetic age by the binary sequel of methylated and non-methylated CpGs on individual DNA strands, which might also be used as a surrogate for the heterogeneity of epigenetic age within a sample.

### Genetic background impacts on epigenetic age-predictions of mice

We have previously demonstrated, that epigenetic age-predictions with our 3 CpG pyrosequencing age-predictor are accelerated in DBA/2 mice, as compared to C57BL/6 mice, which may reflect the different life expectancy of these mouse strains (Han et al., 2018). Furthermore, we demonstrated that age-predictions with this predictor were also accelerated in C57BL/6 mice with quantitative trait locus insertion from DBA/2 into the congenic C57BL/6 chromosome 11, which was expected to be associated with the shorter lifespan of DBA/2 (referred to as Line A mice) (Brown et al., 2019). We now determined, within the same samples, whether the epigenetic age-acceleration can also be observed in DBA/2 mice (n = 33) and Line A mice (n = 15) using the BBA-seq approach. In fact, the predictions with either the 3 CpG BBA-seq, or the 7 CpG BBA-seq Lasso-regression model, provided very similar results as previously observed for the 3 CpG pyrosequencing clock (Figure 5a and b).

**Figure 5.**
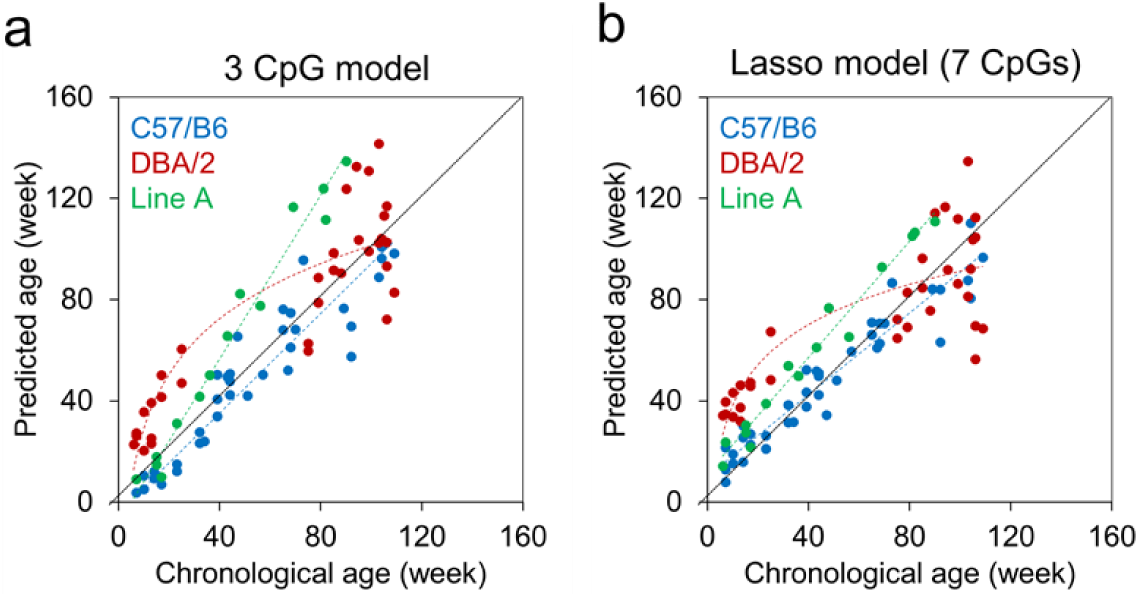
Age-predictions with BBA-seq in mice from different genetic background. DNAm levels were analyzed with BBA-seq in blood samples of 40 C57BL/6 mice of the validation sets, 33 DBA/2 mice, and 15 transgenic C57BL/6 mice with an age-associated region from DBA/2 mice (Line A) (Brown et al., 2019). For epigenetic age predictions we either used (**a**) the 3 CpG multivariable model, or (**b**) the lasso regression model based on 7 CpGs of the same three amplicons (*Prima1, Hsf4, Kcns1)*. As previously described for pyrosequencing, epigenetic age-predictions were logarithmically accelerated in DBA/2 mice (Han et al., 2018), and also accelerated in Line A mice (Brown et al., 2019).

Subsequently, we analyzed the single read patterns of BBA-seq data in DBA/2 and Line A mice. We observed the same random gain or loss of DNAm at neighboring CpGs (Figure 6a) and a moderate correlation in DNAm at neighboring CpGs (Figure 6b), as previously observed for C57BL/6 mice. Furthermore, single read predictions within the three amplicons for *Prima1, Hsf4* and *Kcns1* (based on the training set of C57BL/6 mice) provided similar heterogeneity and acceleration of age-estimations (Figure 6c and d). These results indicate that epigenetic aging is generally accelerated within the three age-associated regions in DBA/2 and Line A mice, as compared to C57BL/6 mice.

**Figure 6.**
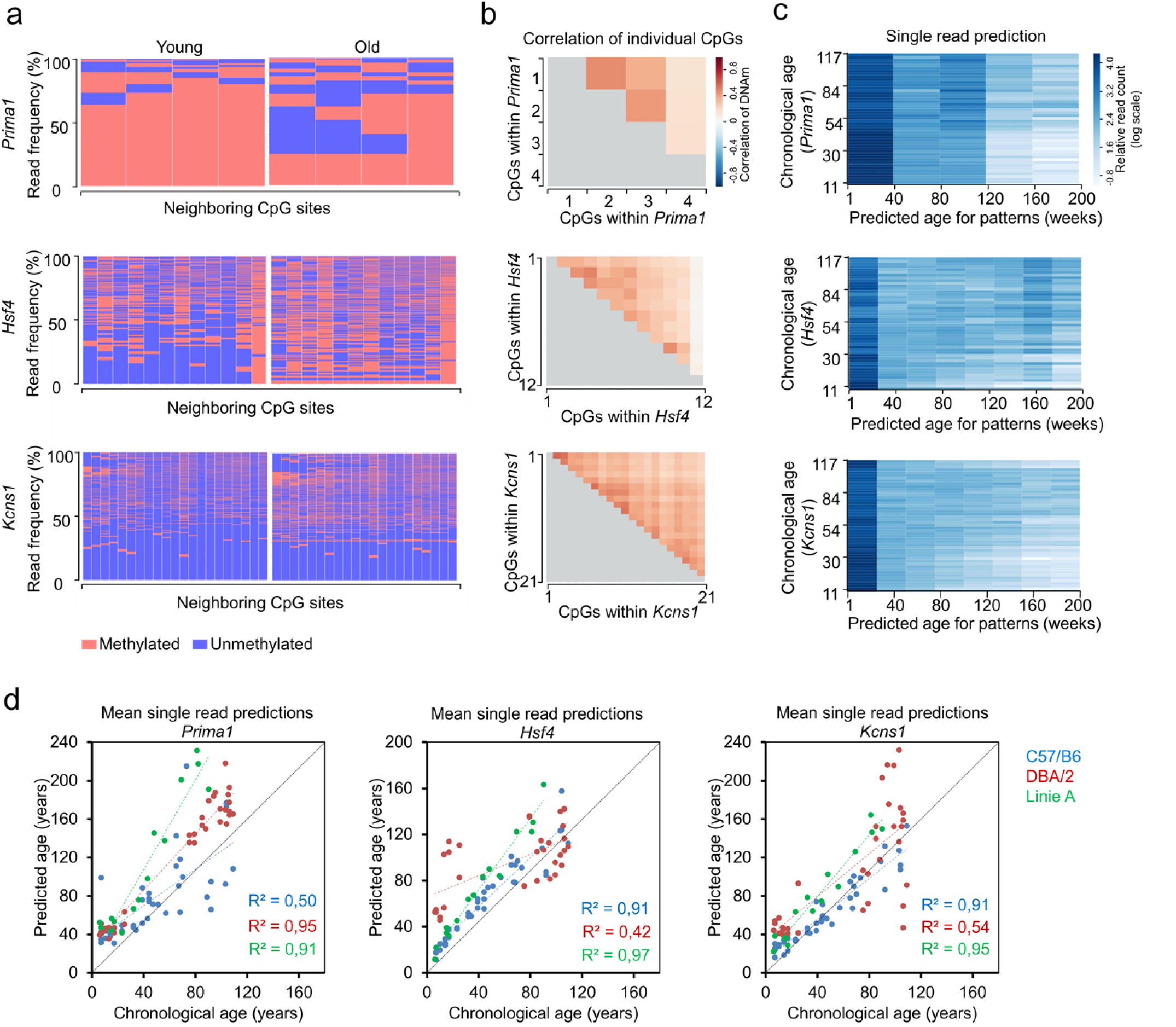
Analysis of age-associated DNAm patterns within individual BBA-seq reads in mice from different genetic background. (**a**) The plots exemplarily display the frequency of DNAm patterns in two DBA/2 mice: one young (7 weeks) and one old DBA/2 mouse(109 weeks). The frequencies of patterns within the amplicons of *Prima1, Hsf4* and *Kcns1* were compared, in analogy to Figure 4a. (**b**) Pearson correlation of DNAm among neighboring CpGs within three amplicons from DBA/2 mice (n = 33). (**c**) Heatmaps of epigenetic age-predictions for individual BBA-seq reads of DBA/2 mice (n = 33). Epigenetic ages were estimated based on the binary sequel of methylated and non-methylated CpGs for three amplicons (read counts were normalized by the readcounts per sample and are depicted in log scale). In analogy to Figure 4c, each read was classified to predicted ages between 0 and 200 weeks. (**d**) The mean of the single read predictions of BBA-seq data was determined for each sample and then plotted against the chronological age of the DBA/2 (n = 33) and line A (n = 15) mice in comparison with validation sets of C57BL/6 mice (n = 40). The linear coefficients of determination (R^2^) of DNAm *versus* chronological age are indicated.

## Discussion

Epigenetic clocks are used as a surrogate marker for the process of biological aging. They are therefore valuable tools to gain insight into effects of aging or rejuvenating interventions (Bell et al., 2019). To this end, the murine model system enables much better standardization over the life-time than what is achievable in humans (Field et al., 2018). The targeted assays for epigenetic clocks are easier applicable than epigenetic clocks that are based on genome wide RRBS or WGBS profiles. A bottleneck for the latter is often a low coverage of reads for specific CpG sites. Particularly pyrosequencing and ddPCR seem to provide more precise measures for DNAm levels at individual CpGs (Blueprint-consortium, 2016). Furthermore, targeted analysis of DNAm only at specific CpGs is faster, facilitates better standardization of procedures, and it is more cost-effective than genome-wide approaches (Wagner, 2017). Thus, the targeted assays may be particularly advantageous for larger intervention studies. On the other hand, the number of CpGs to be implemented into epigenetic clocks provides a tradeoff between accuracy, which is generally increased with more age-associated CpGs, *versus* applicability and costs. In this regard, larger signatures that are based on genome wide DNAm profiles may be advantageous.

It is not trivial to directly compare the performance of our targeted epigenetic clocks with the other published predictors for WGBS or RRBS data, since the tissues, age-ranges, and methods vary considerably in these studies (Meer, Podolskiy, Tyshkovskiy, & Gladyshev, 2018; Petkovich et al., 2017; Stubbs et al., 2017; Wang et al., 2017). The previously published RRBS and WGBS clocks revealed high precision in the training sets, which markedly decreased when tested on independent samples. For murine blood samples, the blood clock by Petkovich et al. showed the best performance with MAE (mean absolute error) of 8.6 weeks (Meer et al., 2018). Our targeted approaches provided similar or sometimes even slightly better accuracy, with an MAE ranging from 4.6 to 12 weeks (or Median error 4.9 to 10 weeks).

In our previous work, we demonstrated that robust and reliable epigenetic age-predictions can be achieved by pyrosequencing at three CpGs (Han et al., 2018). We anticipated that the very high correlation of a 15 CpG lasso regression model, which was suggested during the review process, might be due to overfitting with the relatively small training set (Han et al., 2018). In the current study, we revisited this model to demonstrate that it indeed provides higher accuracy and precision than the 3 CpG predictor - however, it also necessitates pyrosequencing of two additional amplicons. It therefore depends on the experimental design and resources which of the pyrosequencing clocks is better suited.

Upon bisulfite conversion, there is a difference in the sequence of methylated and non-methylated DNA and this can entail a PCR bias (Warnecke et al., 1997). Such DNAm sensitive PCR bias might be reduced by ddPCR, since it relies on detection of either methylated or non-methylated DNA in individual droplets, rather than the amplification efficiency (Weisenberger et al., 2008). So far, ddPCR is particularly applied for detection and quantification of genetic aberrations. Several studies demonstrated that it also enables precise measurements of DNAm levels (Hindson et al., 2013; Yu, Heinzerling, & Grady, 2018; Zemmour et al., 2018), while only few recent studies reported ddPCR assays for epigenetic clocks in humans (Han et al., 2020; Shi et al., 2018). A major challenge for the establishment of such assays is the design of reliable and specific primers and probes for the bisulfited converted DNA sequences. In this study, we describe a 3 CpG ddPCR assay, that facilitates similar accuracy in age-predictions as the previously described 3 CpG pyrosequencing assay.

Next generation sequencing platforms enable targeted DNAm analysis in a barcoded manner for multiple samples in parallel (Han et al., 2020; Naue et al., 2018). In this study, we describe that BBA-seq of only three age-associated regions facilitates also reliable epigenetic age-predictions for murine blood samples. Advantages of this approach are the very high coverage and the relatively long target regions (up to 500 base pairs), which may cover more neighboring CpGs than pyrosequencing or ddPCR (Franzen et al., 2017). Our results confirmed that the correlation of chronological age with DNAm levels follows a bell-shaped curve at neighboring CpGs within about 200 to 400 bases of BBA-seq amplicons (Han et al., 2020) – this was particularly observed in amplicons of *Hsf4* and *Kcns1* that comprised more neighboring CpGs. On the other hand, within individual BBA-seq reads there was only a moderate correlation of DNAm at neighboring CpGs. This is further substantiated by the mean single read predictions which clearly correlate with chronological age. Thus, our results support the notion that age-associated genomic regions favor a stochastic accumulation of DNAm changes, which may be attributed to other epigenetic modifications or higher chromatin order. If age-associated DNAm was directly medicated by epigenetic writers, such as DNMTs or TETs, it might be anticipated that neighboring CpGs are rather coherently modified. The functional relevance of these age-associated DNAm changes remains unclear. Altered promoter methylation with aging was found to be generally unrelated to altered gene expression, also in mice (Hadad, Masser, Blanco-Berdugo, Stanford, & Freeman, 2019). There is evidence, that the epigenetic drift by stochastic DNAm changes in promoters results in degradation of coherent transcriptional networks during aging (Hernando-Herraez et al., 2019). In the future, it will be important to better understand and validate how heterogeneity in single BBA-seq read predictions reflects heterogeneity of epigenetic aging within a sample. To this end, it will be interesting to further investigate single-cell DNAm profiles, longer reads that cover multiple age-associated domains (e.g. by nanopore sequencing), or analysis of single-cell derived clones.

Various epigenetic clocks for mice were demonstrated to reflect aspects of biological aging, rather than only chronological aging (Meer et al., 2018). It is still not unequivocally proven if specific epigenetic clocks capture such aspects of biological aging better, or if they may rather be influenced by the cellular composition or by direct association of DNAm at individual CpGs with specific diseases. We have recently demonstrated that inhibition of Cdc42 activity extends lifespan in C57BL/6 mice, and this is also reflected by younger age-predictions with our 3 CpG pyrosequencing signature (Florian et al., Accepted for publication). In this study, we validated that the shorter-lived DBA/2 mice and the Line A mice have also accelerated epigenetic aging in BBA-seq data – in the conventional epigenetic predictors based on DNAm levels as well as in the single-read BBA-seq predictions for all three amplicons. Thus, our 3 CpG signature clearly captures aspects of biological aging in mice. Furthermore, the accelerated epigenetic aging in DBA/2 and Line A mice cannot be attribute to deviations at individual CpGs, but rather affects the entire age-associated region.

Taken together, we further developed and compared targeted epigenetic clocks for mice with pyrosequencing, ddPCR, or BBA-seq. All three methods provided reliable age-predictions with similar accuracy as previously described for RRBS and WGBS clocks. For DNAm levels at individual CpGs the measurements with pyrosequencing and ddPCR seemed to correlate slightly better with chronological age than BBA-seq results. On the other hand, the longer reads of BBA-seq gave better insight into neighboring CpGs and facilitate even single-read predictions that may reveal heterogeneity in epigenetic aging within a sample – depending on the availability of instruments and the experimental design all of these methods may now be considered for targeted epigenetic clocks in mice.

## Materials and Methods

### Mouse strains and blood collection

Blood specimens of C57BL/6J mice of the training set (n = 24) and of the validation set 1 (n = 21), DBA/2J mice (n = 33), and Line A mice (n = 15) were obtained by submandibular bleeding (100-200 μl) of living mice or postmortem from the vena cava at the University of Ulm. One sample from the training set was excluded in the subsequent ddPCR and BBA-seq analysis due to the lack of bisulfite converted DNA. C57BL/6J samples of the validation set 2 (n = 19) were collected at the University of Groningen from the cheek. All mice were fed by *ad libitum*, and housed under pathogen-free conditions. Experiments were performed in compliance with the Institutional Animal Care of the Ulm University as well as by Regierungspräsidium Tübingen and with the Institutional Animal Care and Use Committee of the University of Groningen (IACUC-RUG), respectively.

### Genomic DNA isolation and bisulfite conversion

Genomic DNA from 50 µl murine blood was isolated by the QIAamp DNA Mini Kit (Qiagen, Hilden, Germany) according to the manufacturer’s instructions. Then, DNA was quantified by Nanodrop 2000 Spectrophotometers (Thermo Scientific, Wilmington, USA). 200 ng of extracted genomic DNA was subsequently bisulfite-converted with the EZ DNA Methylation Kit (Zymo Research, Irvine, USA).

### Pyrosequencing

Bisulfite converted DNA was initially subjected to PCR amplification. Primers were purchased at Metabion and the sequences are provided in Table S1, as described before (Han et al., 2018). 20 µl PCR products were subsequently immobilized to 5 µl Streptavidin Sepharose High Performance Bead (GE Healthcare, Piscataway, NJ, USA), and then were finally annealed to 1 µl sequencing primer (5 μM) for 2 minutes at 80°C. Amplicons were sequenced using PyroMark Gold Q96 Reagents (Qiagen) on PyroMark Q96 ID System (Qiagen, Hilden, Germany) and analyzed with PyroMark Q CpG software (Qiagen). The relevant sequences are depicted for the five relevant genomic regions in Figure S4. The 15 CpG model for pyrosequencing data, which was trained by lasso regression with the lambda parameter chosen by cross-fold validation, has been described before (Han et al., 2018) and is provided in Table S2.

### Droplet digital PCR (ddPCR)

DNA methylation analysis by ddPCR was performed with a QX200TM Droplet Digital™ PCR System (Bio-Rad, CA, USA). We used dual-labeled TaqMan hydrolysis probes which recognize either the methylated or non-methylated target CpG site. All the primers and probes were designed by Primer3Plus software (Table S3). Each 20 μl reaction mixture consisted of 10 μl of 2X ddPCR Supermix (No dUTP; Bio-Rad), 1 μM of the forward and reverse primers, 250 nM of the dual probes, and 25 ng of bisulfite converted DNA. The mixture and 70 μl of droplet generation oil was then subjected into QX200 Droplet Generator (Bio-Rad). 40 μl of the generated droplets were transferred to the ddPCR 96 well plate (Bio-Rad). The plate was heat sealed with the PX1 PCR Plate Sealer (Bio-Rad) and subsequently placed in the C1000 Touch Thermal Cycler (Bio-Rad) for PCR runs as follows: 95°C for 10 min, 40 cycles of 94°C for 30 s and 1 min (2.5°C/s ramp rate) at 55°C (*Prima1, Kcns1*) or 58°C (*Hsf4*), followed by 10 min enzyme deactivation step at 98°C and a final hold at 4°C. The PCR plate was read on the QX200 droplet reader (Bio-Rad) and data were analyzed by QuantaSoft 1.7.4 software (Bio-Rad). The percentage methylation of each reaction was determined by Poisson statistics according to the fraction of positive droplets for methylated and non-methylated probes. The multivariable regression model for ddPCR is provided in Table S4.

### Barcoded bisulfite amplicon sequencing (BBA-seq)

Target sequences (Figure S5) for *Prima1, Hsf4* and *Kcns1* were amplified by PyroMark PCR kit (Qiagen) using forward and reverse primers containing handle sequences for the subsequent barcoding step (Table S5). PCR was run under the following conditions: 95°C for 15 min; 40 cycles of 94°C for 30 s, 60°C for 30 s, 72°C for 30 s; and final elongation 72°C for 10 min. The three amplicons of each donor were pooled at equal concentrations under the quantification of Qubit (Invitrogen), and cleaned up with paramagnetic beads from Agencourt AMPure XP PCR Purification system (Beckman Coulter). 4 μl of pooled products were subsequently added to 21 μl PyroMark Master Mix (Qiagen) containing 10 pmol of barcoded primers (adapted from NEXTflexTM 16S V1-V3 Amplicon Seq Kit, Bioo Scientific, Austin, USA) for a second amplification (95°C for 15 min; 16 cycles of 95°C for 30 s, 60°C for 30s, 72°C for 30s; final elongation 72°C for 10min). PCR products were again quantified by Qubit (Invitrogen), equimolarly pooled, and cleaned up by Select-a-Size DNA Clean & Concentrator Kit (Zymo Research, USA). 10 pM DNA library was prepared under Denature and Dilute Libraries Guide of Illumine MiSeq System with 15% PhiX spike-in control (Illumina, CA, USA) and eventually subjected to 250 bp pair-end sequencing on a MiSeq lane (Illumina, CA, USA) using Miseq reagent V2 Nano kit (Illumina). We utilized the Bismark tool (Krueger & Andrews, 2011) to determine the DNAm levels for each CpG based on BBA-seq data. Multivariable regression models for epigenetic age predictions were generated based on three CpGs that revealed highest correlation with chronological age per amplicon (Table S6). Alternatively, we used a penalized regression model from the R package glmnet on the training dataset to establish a predictor with machine learning (Table S7). The alpha parameter of glmnet was set to 1 (lasso regression) and the lambda parameter was chosen by cross-fold validation of the training dataset (10-fold cross validation).

### Epigenetic age predictions for individual BBA-seq reads

As previously described, we developed an algorithm to estimate epigenetic age based on the binary sequel of methylated and non-methylated CpGs within individual reads of BBA-seq data (Han et al., 2020). In brief, according the age-associated correlations at individual CpG of the BBA-seq training set, each DNAm pattern with binary sequel of methylation and unmethylation was assigned to their most representative corresponding age (0 to 200 weeks). For each donor, we calculated the mean of strand-specific age-predictions weighted by read counts as final epigenetic age predictions. Further details on the rational and derivation of the mathematical model are provided in our previous work (Han et al., 2020).

## Acknowledgement

This work was supported by the German Research Foundation (DFG; WA 1706/8-1 and WA 1706/12-1 to WW), by the German Ministry of Education and Research (BMBF; Epi-Blood-Count to WW; and SyStarR to HG), and by the NIH (R01HL134617 and R01DK104814 to HG). The Groningen samples were obtained from the Mouse Clinic for Cancer and Ageing (http://www.mccanet.nl), which is supported by a grant from the Netherlands Organization for Scientific Research (NWO). The funding bodies were not involved in study design, data analysis, or writing of the manuscript.

## Authors contributions

Yang Han: performed pyrosequencing and BBA-seq, formal analysis, writing of original draft; Miloš Nikolić: performed BBA-seq data analysis; Michael Gobs: performed ddPCR assays; Julia Franzen: calculated alternative aging models; Gerald de Haan: resources and project administration; Hartmut Geiger: resources, funding acquisition; Wolfgang Wagner: conceptualization, supervision, funding acquisition, writing of original draft, project administration.

## References

Bell, C. G., Lowe, R., Adams, P. D., Baccarelli, A. A., Beck, S., Bell, J. T., … Horvath, S. (2019). DNA methylation aging clocks: challenges and recommendations. Genome biology, 20(1), 249.

Belsky, D. W., Caspi, A., Arseneault, L., Baccarelli, A., Corcoran, D. L., Gao, X., … Houts, R. (2020). Quantification of the pace of biological aging in humans through a blood test, the DunedinPoAm DNA methylation algorithm. eLife, 9.

Blueprint-consortium. (2016). Quantitative comparison of DNA methylation assays for biomarker development and clinical applications. Nat Biotechnol, 34(7), 726–737. doi: 10.1038/nbt.3605

Bocklandt, S., Lin, W., Sehl, M. E., Sánchez, F. J., Sinsheimer, J. S., Horvath, S., & Vilain, E. (2011). Epigenetic predictor of age. PLoS One, 6(6), e14821.

Brown, A., Schuetz, D., Han, Y., Daria, D., Nattamai, K. J., Eiwen, K., … van Zant, G. (2019). The lifespan quantitative trait locus gene Securin controls hematopoietic progenitor cell function. Haematologica, haematol. 2018.213009.

Dor, Y., & Cedar, H. (2018). Principles of DNA methylation and their implications for biology and medicine. The Lancet, 392(10149), 777–786.

Fahy, G. M., Brooke, R. T., Watson, J. P., Good, Z., Vasanawala, S. S., Maecker, H., … Horvath, S. (2019). Reversal of epigenetic aging and immunosenescent trends in humans. Aging cell, 18(6), e13028.

Field, A. E., Robertson, N. A., Wang, T., Havas, A., Ideker, T., & Adams, P. D. (2018). DNA methylation clocks in aging: categories, causes, and consequences. Molecular cell, 71(6), 882–895.

Florian, M., Leins, H., Gobs, M., Han, Y., Marka, G., Soller, K., … Geiger, H. (Accepted for publication). Inhibition of Cdc42 activity extends lifespan and decreases circulating inflammatory cytokines in aged female C57BL/6 mice. Aging cell.

Franzen, J., Zirkel, A., Blake, J., Rath, B., Benes, V., Papantonis, A., & Wagner, W. (2017). Senescence-associated DNA methylation is stochastically acquired in subpopulations of mesenchymal stem cells. Aging cell, 16(1), 183–191.

Gross, A. M., Jaeger, P. A., Kreisberg, J. F., Licon, K., Jepsen, K. L., Khosroheidari, M., … Ng, C. T. (2016). Methylome-wide analysis of chronic HIV infection reveals five-year increase in biological age and epigenetic targeting of HLA. Molecular cell, 62(2), 157–168.

Hadad, N., Masser, D. R., Blanco-Berdugo, L., Stanford, D. R., & Freeman, W. M. (2019). Early-life DNA methylation profiles are indicative of age-related transcriptome changes. Epigenetics & chromatin, 12(1), 58.

Han, Y., Eipel, M., Franzen, J., Sakk, V., Dethmers-Ausema, B., Yndriago, L., … Wagner, W. (2018). Epigenetic age-predictor for mice based on three CpG sites. eLife, 7, e37462.

Han, Y., Franzen, J., Stiehl, T., Gobs, M., Kuo, C.-C., Nikolić, M., … Ritz-Timme, S. (2020). New targeted approaches for epigenetic age predictions. BMC Biology, 18(1), 1–15.

Hernando-Herraez, I., Evano, B., Stubbs, T., Commere, P.-H., Bonder, M. J., Clark, S., … Reik, W. (2019). Ageing affects DNA methylation drift and transcriptional cell-to-cell variability in mouse muscle stem cells. Nature communications, 10(1), 1–11.

Hindson, C. M., Chevillet, J. R., Briggs, H. A., Gallichotte, E. N., Ruf, I. K., Hindson, B. J., … Tewari, M. (2013). Absolute quantification by droplet digital PCR versus analog real-time PCR. Nature methods, 10(10), 1003–1005.

Horvath, S., Erhart, W., Brosch, M., Ammerpohl, O., von Schönfels, W., Ahrens, M., … Spector, T. D. (2014). Obesity accelerates epigenetic aging of human liver. Proceedings of the National Academy of Sciences, 111(43), 15538–15543.

Horvath, S., Garagnani, P., Bacalini, M. G., Pirazzini, C., Salvioli, S., Gentilini, D., … Vinters, H. V. (2015). Accelerated epigenetic aging in Down syndrome. Aging cell, 14(3), 491–495.

Horvath, S., & Raj, K. (2018). DNA methylation-based biomarkers and the epigenetic clock theory of ageing. Nature Reviews Genetics, 1.

Koch, C. M., & Wagner, W. (2011). Epigenetic-aging-signature to determine age in different tissues. Aging (Albany NY), 3(10), 1018–1027. doi:100395 [pii]

Krueger, F., & Andrews, S. R. (2011). Bismark: a flexible aligner and methylation caller for Bisulfite-Seq applications. bioinformatics, 27(11), 1571–1572.

Maegawa, S., Lu, Y., Tahara, T., Lee, J. T., Madzo, J., Liang, S., … Issa, J.-P. J. (2017). Caloric restriction delays age-related methylation drift. Nature communications, 8(1), 539.

Maierhofer, A., Flunkert, J., Oshima, J., Martin, G. M., Haaf, T., & Horvath, S. (2017). Accelerated epigenetic aging in Werner syndrome. Aging (Albany NY), 9(4), 1143.

Marioni, R. E., Shah, S., McRae, A. F., Chen, B. H., Colicino, E., Harris, S. E., … Cox, S. R. (2015). DNA methylation age of blood predicts all-cause mortality in later life. Genome biology, 16(1), 25.

Meer, M. V., Podolskiy, D. I., Tyshkovskiy, A., & Gladyshev, V. N. (2018). A whole lifespan mouse multi-tissue DNA methylation clock. eLife, 7, e40675.

Naue, J., Sänger, T., Hoefsloot, H. C., Lutz-Bonengel, S., Kloosterman, A. D., & Verschure, P. J. (2018). Proof of concept study of age-dependent DNA methylation markers across different tissues by massive parallel sequencing. Forensic Science International: Genetics, 36, 152–159.

Petkovich, D. A., Podolskiy, D. I., Lobanov, A. V., Lee, S.-G., Miller, R. A., & Gladyshev, V. N. (2017). Using DNA methylation profiling to evaluate biological age and longevity interventions. Cell Metabolism, 25(4), 954-960. e956.

Shi, L., Jiang, F., Ouyang, F., Zhang, J., Wang, Z., & Shen, X. (2018). DNA methylation markers in combination with skeletal and dental ages to improve age estimation in children. Forensic Science International: Genetics, 33, 1–9.

Stubbs, T. M., Bonder, M. J., Stark, A.-K., Krueger, F., von Meyenn, F., Stegle, O., & Reik, W. (2017). Multi-tissue DNA methylation age predictor in mouse. Genome biology, 18(1), 68.

Wagner, W. (2017). Epigenetic aging clocks in mice and men. Genome biology, 18(1), 107.

Wang, T., Tsui, B., Kreisberg, J. F., Robertson, N. A., Gross, A. M., Yu, M. K., … Ideker, T. (2017). Epigenetic aging signatures in mice livers are slowed by dwarfism, calorie restriction and rapamycin treatment. Genome biology, 18(1), 57.

Warnecke, P. M., Stirzaker, C., Melki, J. R., Millar, D. S., Paul, C. L., & Clark, S. J. (1997). Detection and measurement of PCR bias in quantitative methylation analysis of bisulphite-treated DNA. Nucleic acids research, 25(21), 4422–4426.

Weisenberger, D. J., Trinh, B. N., Campan, M., Sharma, S., Long, T. I., Ananthnarayan, S., … McCormick, F. (2008). DNA methylation analysis by digital bisulfite genomic sequencing and digital MethyLight. Nucleic acids research, 36(14), 4689–4698.

Yu, M., Heinzerling, T. J., & Grady, W. M. (2018). DNA Methylation Analysis Using Droplet Digital PCR. Methods Mol Biol, 1768, 363–383.

Zemmour, H., Planer, D., Magenheim, J., Moss, J., Neiman, D., Gilon, D., … Landesberg, G. (2018). Non-invasive detection of human cardiomyocyte death using methylation patterns of circulating DNA. Nature communications, 9(1), 1443.

