## Supplementary material for "Epigenetic age-predictions in mice using pyrosequencing, droplet digital PCR or barcoded bisulfite amplicon sequencing": Figures S1-S5 and Tables S1-S7

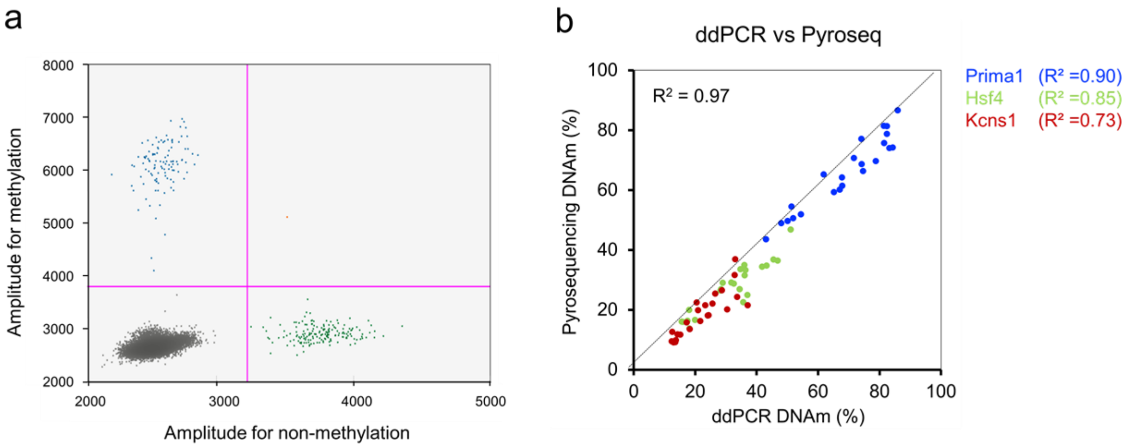


**Figure S1. Comparative analysis of age-associated DNAm by ddPCR and Pyrosequencing.**
(**a**)This plot exemplarily depicts ddPCR results of DNAm in *Prima1* (Blue: positive droplets for methylated; Green: positive droplets for unmethylated; Orange: double positive droplets; Grey: negative droplets). (**b**) DNAm analysis at three age-associated regions were compared in 23 samples with ddPCR and pyrosequencing. Overall, the DNAm levels correlated very good between the two methods.

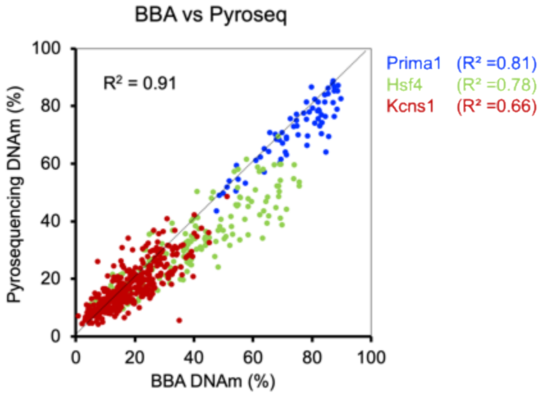


**Figure S2. Comparison of DNAm levels in pyrosequencing and BBA-seq.**
The DNAm levels at the three age-associated genomic regions were compared in 23 blood samples with barcoded bisulfite amplicon sequencing (BBA-seq) and pyrosequencing. All CpGs that were covered by both types of measurements were included into the comparison.

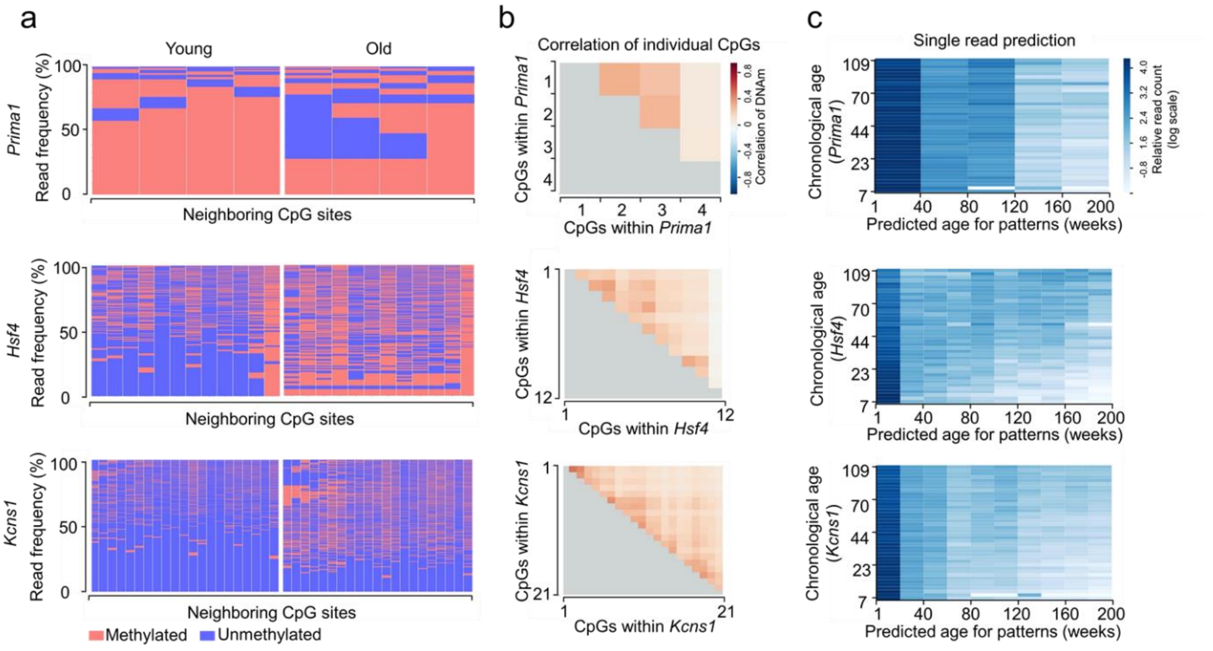


**Figure S3. Analysis of age-associated DNAm patterns within individual BBA-seq reads in C57BL/6 mice of the validation sets.**

(**a**) Heatmaps display the frequency of DNAm patterns within *Prima1*, *Hsf4* and *Kcns1* amplicons from BBA-seq data. One young (7 weeks) and one old (109 weeks) C57BL/6 mouse samples from BBA-seq validation sets were compared. (**b**) Pearson correlation of DNAm among neighboring CpGs within three amplicons from BBA-seq validation sets. (**c**) According to single BBA-seq read prediction of the training set, epigenetic ages were estimated based on the binary sequel of methylated and non-methylated CpGs for three amplicons separately. Each plot classified relative read count of each donor in the two validation sets for predicted ages between 0 and 200 weeks (Relative read count was normalized by logarithmic scale of read count per sample).

*Prima1*

AATTATGAATTAGGTTTATATTTTTCGGGTGGGGGGTAACGA**CG**GTTTT

Sequencing primer → 1 2 3

*Hsf4*

ATTGTAGAAAAAATGGGAAGGTATTAATGTTGGTATTTTTGGTTTTGTTTTATGTGTTTCGGATGGTGTTTTTTGTTTGTAG

Sequencing primer → 1

GTATTTGCGTTG**CG**AGG**CG**ATGATAGT**CG**ATGGCGTT**CG**GAAGA

2 3 4 5 6 7

*Kcns1*

GATATTTCGAAGTTGAATTAAG**CG**ATGTAGAAGTATTTTAGGCGG**CG**TAGTAT**CG**GGT**CG**T**CG**CGTATTTTTTTTGCGTTG

Sequencing primer → 1 2 3 4 5 6 7 8

CGATTCGCGGTTATTGTAGTTATCGT**CG**T**CG**TCGTTTTTCGAGTTTG

9 10 11 12 13 14 15 16

*Gm9312*

GGT**CG**TCGCGGTGTTTCGGTATTGGGTCGAG**CG**CGTGGTGAAATTAGAGGTCGTGGGCGTT

7 6 5 4 3 2 1 ← Sequencing primer

*Gm7325*

TTTATGTTTTGGGAGTTTAGTCGGCGGGTTAGT**CG**GCGTAG

Sequencing primer → 1 2 3 4

**Figure S4. Targeted sequences of pyrosequencing assays**.
Sequences for the five genomic regions (after bisulfite conversion). The CpG sites are depicted in red and numbered consecutively by their dispensation order. The relevant 15 CpGs selected by the machine learning model are highlighted in bold.

**1. *Prima1* (chr12:103,214,592-103,214,742)**

CTGTGCCTAATTAGGAGAGGCAAATTATGAATCAGGTTCACATCTCT**CG**GGTGGGGGGCAACGACGGCTTTTCCTGGGTGAGGAAAAGATCACAG

48

GGAGGTCTGAGCAAGAGC**CG**GGAACAGCAGGTCACAAGCTGGTGTAATTAATTTTG

65

**2. *Hsf4* (chr8:105,270,895-105,271,241)**

AGGTGGGACAAACTGCAGAAAAAATGGGAAGGCACCAATGTTGGTACTTCTGGCTCTGCCTTATGTGTCCCGGATGGTGCCTCCTGCCTGCAGGTA

71

CCTGCGCTGCGAGGCGATGACAGTCGATGG**CG**TCCGGAAGACCTGAGCCGACTGCTGGGAGAGGTGCAAGCTTTGAGAGGAGTGCAGGAGAGC

127

ACGGAGGCACGGCTGCAGGAACTCAGGCAGTGCGGGACAGGGAAGGAAGGGTGGAGTTAGGGGTGAGAGGCTCACTGGCCACCTAGGGGTTC

191

TGGGCAGCTCCTTCCGCTCTTTGGTTGGAGCCTCCATGAAGACAATGGGTCCAACTCTGAAACTTT

**3. *Kcns1* (chr2:164,168,030-164,168,240)**

AGAAGTTGCGCGTGCTGGGAGCCAGCAGCAGG**CG**CGACGACACCTCGAAGCTGAACCAAGCGATGCAGAAGTACTCCAGGCGGCGCAGCACCG

33

GGTCGTCGCGCACCTCCTCTG**CG**CTG**CG**ACCCGCGGCCACTGCAGCCACCGCCGC**CG**CCGCCTCCCGAGCTTGGTACTCCGGCAGGCTGTGGATG

115 120 149

CACATGGCAGCGATGGAGGCGAG

**Figure S5. Targeted sequences for BBA-seq.**
Sequences for nine genomic regions that were analyzed by BBA-seq are depicted and all CpG sites are highlighted in red. The age-related CpGs selected by machine learning approach are highlighted in bold and by their ordering number in the sequences (underlined CpG were used in the multivariable linear model).

**Table S1. Primers for pyrosequencing assays.**

| Primer | Sequence |
|---|---|
| ***Prima1*** | |
| Forward | 5'-TTGTGTTTAATTAGGAGAGGTAAATTATGAATTAGGTTTATA-3' |
| Reverse | 5'-Biotin-CAAAATTAATTACACCAACTTATAACCTACTATTC-3' |
| Sequencing | 5'-AATTATGAATTAGGTTTATATTT-3' |
| ***Hsf4*** | |
| Forward | 5'-GTGAGTAGTAAGGTGGGATAAATTGTAGAAAAAATG-3' |
| Reverse | 5'-Biotin-TCCCTACTCTCCTACACTCCTCTCAAAACTTA-3' |
| Sequencing | 5'-ATTGTAGAAAAAATGGGAA-3' |
| ***Kcns1*** | |
| Forward | 5'-GGTTGAGAGGGTGGTAGAAGAAGTTG-3' |
| Reverse | 5'-Biotin-ACTCCCCTCCATCCCTACCATATACATCCA-3' |
| Sequencing | 5'-GAAGATATTTAGAAGTTGAATT-3' |
| ***Gm9312*** | |
| Forward | 5'-Biotin-TTGTTTTGGGGTATTAGAAATTTTTTT-3' |
| Reverse | 5'-CCTAACCATACTAAACCAAATCTCTATATCTAAAT-3' |
| Sequencing | 5'-AACCCCCACCACCTCTAATTTCAC-3' |
| ***Gm7325*** | |
| Forward | 5'-TGTTGGTTGAGGATAAAGAGTAGATAGTTTAGTAGAGT-3' |
| Reverse | 5'-Biotin-TTCCCTTTACAAATACAAATCCTACCATA-3' |
| Sequencing | 5'-TTTATGTTTTGGGAGTTTA-3' |

**Table S2. Machine learning model based on 15 CpGs (Pyrosequencing)**

| Gene name | Chromosome | CpG# | Position | $R^2$ | Coefficients |
|---|---|---|---|---|---|
| (Intercept) | | | | | -1.62345 |
| *Prima1* | chr12 | 3 | 103214656 | 0,62 | -0.29869 |
| *Hsf4* | chr8 | 3 | 105271000 | 0,95 | 1.254757 |
| *Hsf4* | chr8 | 4 | 105271006 | 0,91 | 0.327821 |
| *Hsf4* | chr8 | 5 | 105271015 | 0,90 | 0.467297 |
| *Hsf4* | chr8 | 7 | 105271025 | 0,81 | 0.001369 |
| *Kcns1* | chr2 | 1 | 164168088 | 0,81 | 0.170725 |
| *Kcns1* | chr2 | 3 | 164168113 | 0,84 | 0.089469 |
| *Kcns1* | chr2 | 4 | 164168121 | 0,74 | 0.320572 |
| *Kcns1* | chr2 | 5 | 164168126 | 0,73 | 0.343513 |
| *Kcns1* | chr2 | 6 | 164168129 | 0,79 | 0.05701 |
| *Kcns1* | chr2 | 13 | 164168175 | 0,82 | 0.326642 |
| *Kcns1* | chr2 | 14 | 164168178 | 0,81 | 0.307202 |
| *Gm9312* | chr12 | 2 | 24252014 | 0,47 | 0.055652 |
| *Gm9312* | chr12 | 7 | 24252050 | 0,25 | 0.434457 |
| *Gm7325* | chr17 | 3 | 45601568 | 0,37 | -0.13861 |

This model was trained with a machine learning approach (linear model with L1 regularization; lambda = 1.570242), as described before (Han et al., 2018). CpG# indicates the relevant age-related CpGs of the amplicon, as also depicted in Figure S4.

**Table S3. Primers for ddPCR assays.**

| Primer | Sequence |
|---|---|
| ***Prima1*** | |
| Forward | 5'- GGAGAGGTAAATTATGAATTAGG -3' |
| Reverse | 5'- ACTCTTACTCAAACCTCCCT -3' |
| Probe | 6-Fam - TATATTTTTCGGGTGGGGG -BHQ-1[a] |
| | Hex- TATATTTTTTGGGTGGGGG -BHQ-1[b] |
| ***Hsf4*** | |
| Forward | 5'- AATGTTGGTATTTTTGGTTTTGTTT -3' |
| Reverse | 5'- AAAACTTACACCTCTCCCAACAAT -3' |
| Probe | 6-Fam - TGTGTTTCGGATGGTGTTTTTTGT - BHQ-1[a] |
| | Hex - TGTGTTTTGGATGGTGTTTTTTGT - BHQ-1[b] |
| ***Kcns1*** | |
| Forward | 5'- TTGGGAGTTAGTAGTAGGYG -3' |
| Reverse | 5'- ATACATCCACAACCTACCRA -3' |
| Probe | 6-Fam - AGTTGAATTAAGCGATGTAGAAGTATTTTA - BHQ-1[a] |
| | Hex - AGTTGAATTAAGTGATGTAGAAGTATTTTA - BHQ-1[b] |

[a] Probe targeting the methylated sequence.
[b] Probe targeting the non-methylated sequence.

**Table S4. Multivariable model for ddPCR.**

| Gene Name | CHR | Map Info | Coefficients | P value |
|---|---|---|---|---|
| (Intercept) | | | -11.56 | 0.83 |
| *Prima1* | 12 | 103214640 | -0.26 | 0.60 |
| *Hsf4* | 8 | 105270966 | 2.33 | 0.001 |
| *Kcns1* | 2 | 164168063 | 0.43 | 0.63 |

**Table S5. Primers for BBA-seq assays.**

| Primer | Sequence |
|---|---|
| ***Prima1*** | |
| Forward | 5'- CTCTTTCCCTACACGACGCTCTTCCGATCTTTGTGTTTAATTAGGAGAGGTAAATTATGAATTAGGTTTATA -3' |
| Reverse | 5'- CTGGAGTTCAGACGTGTGCTCTTCCGATCTCAAAATTAATTACACCAACTTATAACCTACTATTC -3' |
| ***Hsf4*** | |
| Forward | 5'- CTCTTTCCCTACACGACGCTCTTCCGATCTAGGTGGGATAAATTGTAGAAAAAATG -3' |
| Reverse | 5'- CTGGAGTTCAGACGTGTGCTCTTCCGATCTAAAATTTCAAAATTAAACCCATTATCTTCA -3' |
| ***Kcns1*** | |
| Forward | 5'- CTCTTTCCCTACACGACGCTCTTCCGATCTAGAAGTTGYGYGTGTTGGGAGT -3' |
| Reverse | 5'- CTGGAGTTCAGACGTGTGCTCTTCCGATCTCTCRCCTCCATCRCTACCATATAC -3' |

**Table S6. Multivariable model for BBA-seq.**

| Gene Name | CHR | Map Info | Coefficients | P value |
|---|---|---|---|---|
| (Intercept) | | | 11.92 | 0.74 |
| *Prima1*.48 | 12 | 103214640 | -0.53 | 0.15 |
| *Hsf4*.127 | 8 | 105271022 | 1.08 | 0.03 |
| *Kcns1*.33 | 2 | 164168063 | 1.53 | 0.003 |

The numbers at the target sites indicate the relevant age-related CpGs of the amplicon, as also depicted in Figure S5.

**Table S7. Machine learning model by Lasso (BBA-seq)**

| | Target sites* | CHR | Map Info | Coefficients |
|---|---|---|---|---|
| | (Intercept) | | | 5.18 |
| 1 | *Prima1*.65 | 12 | 103214657 | -0.26 |
| 2 | *Hsf4*.71 | 8 | 105270966 | 0.83 |
| 3 | *Hsf4*.191 | 8 | 105271086 | 0.11 |
| 4 | *Kcns1*.33 | 2 | 164168063 | 0.19 |
| 5 | *Kcns1*.115 | 2 | 164168145 | 0.41 |
| 6 | *Kcns1*.120 | 2 | 164168150 | 0.52 |
| 7 | *Kcns1*.149 | 2 | 164168179 | 0.43 |

This model was trained with a machine learning approach, although the number of samples may not be enough for cross-validation on the data. The model should therefore be retained when a larger set of samples is available. The numbers at the target sites indicate the relevant age-related CpGs of the amplicon, as also depicted in Figure S5.
